# Telomere-to-Telomere Assembly Improves Host Reads Removal in Metagenomic High-Throughput Sequencing of Human Samples

**DOI:** 10.1101/2023.05.05.539517

**Authors:** Lei Wang, Guoning Xing

**Affiliations:** Department of Bioinformatics Research, Genskey Co., Ltd., Beijing 102206, China

## Abstract

Metagenomic high-throughput sequencing brings revolution to the study of human microbiome, clinical pathogen detection, discovery and infection diagnosis, but clinical samples often contain abundant human nucleic acids, leading to a high proportion of host reads. A high-quality human reference genome is essential for removing host reads to make downstream analyses faster and more accurate. The recently published complete human genome, Telomere-to-Telomere CHM13 assembly (T2T), though achieved great success immediately, has yet to be tested for metagenomic sequencing. In this study, we demonstrated the innovation that T2T brings to the field, using a diverse set of samples containing 4.97 billion reads sequenced from 165 libraries, on short- and long-read platforms. To exclude the effect of algorithms in comparison of the genomes, we benchmarked the per-read performance of state-of-the-art algorithms. For short reads, bwa mem was the best-performing algorithm, with positive median of differences (MD) and adjusted p-values <0.001 for all comparisons, while no consistent difference in overall performance was found for long reads algorithms. T2T, when compared to current reference genomes hg38 and YH, significantly improved the per-read sensitivity (MD: 0.1443 to 0.7238 percentage point, all adjusted p-values < 0.001) in removing host reads for all sequencers, and the per-read Mathew’s correlation coefficient (MCC) with T2T was also higher (MD: 1.063 to 16.41 percentage point, all adjusted p-values <0.001). Genomic location of reads exclusively mappable to T2T concentrated mainly in newly added regions. Misclassified reads generally resulted from low complexity sequences, contaminations in reference genomes and sequencing abnormalities. In downstream microbe detection procedures, T2T did not affect true positive calls but greatly reduced false positive calls. The improvement in the ability to correctly remove host reads foretells the success of T2T to serve as the next prevailing reference genome in metagenomic sequencing of samples containing human nucleic acids.

---

Metagenomic high-throughput sequencing has been an increasingly valuable tool in rapid pathogen detection and discovery, usually for infection diagnosis in clinical settings. It brought revolution to the field, since it was faster, relatively unbiased, more specific and more sensitive than traditional techniques.^1^ Noteworthily, its application played crucial role in the rapid identification of SARS-CoV-2 in the early stage of the COVID-19 pandemic.^2^ However, most clinical samples contain considerable amount of human nucleic acids, leading to a high proportion of human reads. Therefore, removing host reads is an essential step for fast and accurate microbe identification and other analyses, for processing the uninformative host reads not only consumes time, but also introduces errors. In addition, there may be legal and ethical reasons that make it desirable to remove the human reads sequenced from clinical samples. Currently, all commonly applied methods of host read removal ultimately involve using a human genome as reference. Since its birth^3,4^, the human reference genome assembly, maintained by Genome Reference Consortium, has undergone 5 major updates. The latest update, Build 38 (GRCh38 or hg38)^5^, has been validated clinically to be used in high-throughput sequencing.^6^ Another mainstream reference genome used in many studies^7,8^ is YH, a haplotype-resolved genome assembly sourced from an Asian individual. However, these genomes contain collapsed duplications, missing sequences, inaccurate assemblies and other issues, mainly due to limitations in assembling short reads.^9,10^ These defects result in a loss of accuracy in removing human reads in metagenomic sequencing. Recently, T2T, a complete human genome, was published. It added almost 200Mbp of new sequence, closed all remaining gaps, and corrected thousands of structural errors, thanks to advances in ultralong sequencing and assembly methods.^9,11^ Immediately, T2T has been applied to many areas and opened up opportunities for new discoveries and applications, including epigenetics, genetic variation, comparative genomics, evolutionary genomics, etc.^10,12–14^ In spite of that, the application of T2T as reference genome has not been tested in metagenomic high-throughput sequencing where host reads removal is needed.

Hereby we conducted a benchmarking experiment to test whether switching to the new T2T genome from current reference genomes would bring benefits to the field, and to test the performance of the state-of-the-art bioinformatic algorithms for host reads removal. These include alignment algorithms for short reads and long reads, namely bowtie2^15^, bwa mem^16^, minimap2^17^ and winnowmap^18^, as well as taxonomy labeling algorithms based on k-mer counting, namely kraken2^19^ and the human read removal tool (HRRT)^20^. HRRT, also known as the sra-human-scrubber, is a tool based on Sequence Taxonomic Analysis Tool (STAT) and is used by the National Center for Biotechnology Information (NCBI) Sequence Read Archive (SRA) to remove any human read from a clinical infection sample before it can be safely uploaded for SRA submission.^21^ In order to compare the performance of the genomes and algorithms, we obtained a read set by sequencing a diverse collection of samples and established the “gold standard” for each read by a voting and conflict-resolution approach (details in the Materials and Methods section). The reference genomes and algorithms were then evaluated by their per-read sensitivity, specificity and Mathew’s correlation coefficients (MCC).^22^

## Results

The benchmarking read set consisted of 4.97 billion reads from 165 DNA libraries of diverse origins, including 67 sequenced on Illumina (ILMN) sequencer, 68 on MGI and 30 on ONT. We obtained on average 23.3 million short reads (mean length 73.6) per library for ILMN, 50.1 million short reads (mean length 49.9) for MGI, and 0.224 million long reads (mean length 3886.6) for ONT. The quality of the reads was assessed to be 94.5% (mean q20) for ILMN, 97.5% (mean q20) for MGI and 92.5% (mean q10) for ONT (Supplementary Figure S1).

Per-read “ground truth”, or gold standard, was determined by first collating the results of every method tested and then resolving the discrepancy by BLAST search (Supplementary Figure S2). The collating process for all the 4.97 billion reads yielded on average 55.6% consensus host labels and 16.6% consensus non-host labels, with the rest being multi-labelled (Figure 1A). After resolving the discrepancy in the multi-labelled reads, we established the gold-standard for comparison, resulting in - (83.3% on average) host reads per sample (Figure 1B). We observed a great variance in the host read ratio per sample, which ideally represented the diversity of real-world clinical samples in terms of their host nucleic acids content. All reads were categorized as true positive (TP), false positive (FP), true negative (TN), and false negative (FP), by comparing the labels of each method to the gold standard. Per-read sensitivity, specificity, and MCC were then calculated for all methods. (Supplementary Table S1-S3) Pairwise comparison of MCC between all methods led to the following results.

**Figure 1.**
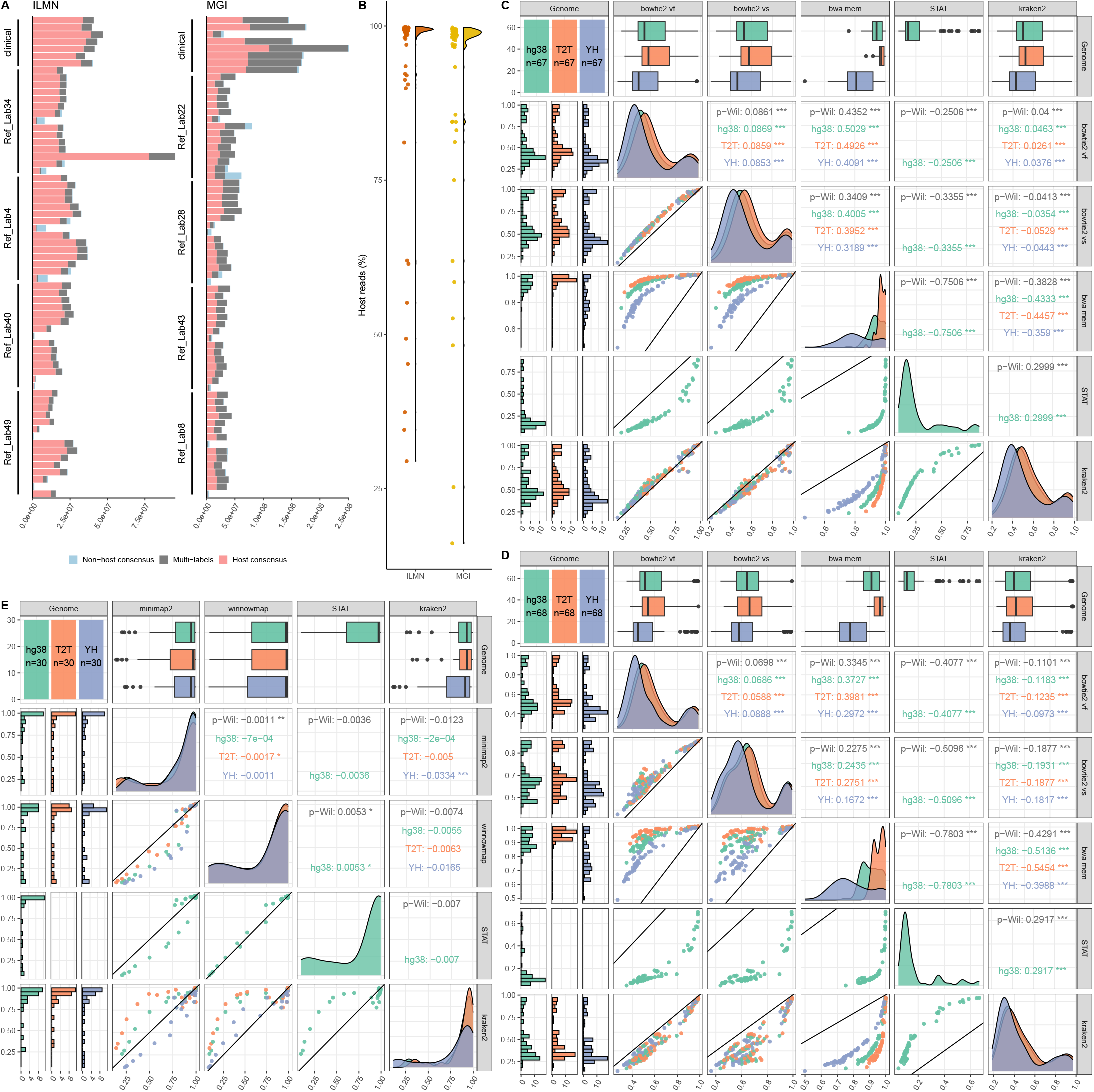
Summary of the performance comparisons. **A** The total number of reads in each sample. Reads are grouped based on the labels given by all host removal methods: non-host consensus (blue), host consensus (red), and multi-labels (grey). **B** Distribution of host reads percentage of all samples, for ILMN (red), MGI (yellow) and ONT (blue). **C-E** Mathew’s correlation coefficient (MCC) comparison across all reference genomes and relevant algorithms. Plots on the diagonal, first column and first row: distribution of per-sample MCC for all genomes, hg38 (green), T2T (orange), and YH (purple). Plots in the upper triangle: paired Wilcoxon (p-Wil) comparisons between each combination of reference genomes and algorithms. Numbers are median of differences between the pair, asterisks show the significance of adjusted p-values (no label: insignificant, *: <0.05, **: <0.01, ***: <0.001), and numbers in colors show the comparison for each genome. Plots in the lower triangle: scatterplot of MCC. The black line has slope of 1 and intercept of 0, therefore points (each representing a sample) above it have higher MCC value in y-axis than x-axis and vice versa. Abbreviations are vf: very fast, vs: very sensitive. Results of different sequencers were arranged in **C** for ILMN, **D** MGI, and **E** ONT.

### The best-performing algorithm for short reads was bwa mem while algorithms for long reads each had its advantages

For short-read sequencing, bwa mem was performing better than others by a large margin, showing significantly higher MCC in all comparisons (omnibus p-value < 0.001, median pairwise differences and *post hoc* p-values in Figure 1C, 1D). Bowtie2, running in very sensitive mode, was the runner-up, followed by kraken2 and bowtie2-in-very-fast-mode, sequentially. The difference between the three were significant, but relatively small (p-values in Figure 1C, 1D). Noteworthily, although less competitive in terms of sensitivity and overall performance, STAT was the most specific algorithm (p-values in Supplementary Figure S3, S4). Being more conservative for short reads made STAT a more desirable option in situations where preserving microbe reads rather than purging host reads was of the uttermost priority.

For long-read sequencing, however, there was no clear winner. Differences in the overall performance of all algorithms, measured by MCC, was statistically insignificant (omnibus p-value 0.6498) (Figure 1E). However, we noticed that kraken2 for long reads was the least specific but most sensitive (p-values in Supplementary Figure S3, S4), therefore performed better on samples richer of human nucleic acids. Nevertheless, it should be noted that in samples which kraken2 outperformed the others, the difference was bigger than *vice versa*, and this was not fully reflected in the nonparametric test for MCC. Therefore, kraken2 was arguably the best performing algorithm for long reads. Sensitivity of minimap2 and winnowmap were slightly lower than that of kraken2 and higher than that of STAT, but in general showed no significant difference to each other (p-values in Supplementary Figure S3). In terms of specificity, no significant difference was found in general between algorithms other than kraken2 (p-values in Supplementary Figure S4).

### T2T improved performance compared to current reference genomes

T2T, out of the three reference genomes, achieved best results in terms of sensitivity and overall performance, for both short- and long-read sequencing. (Figure 1C-E, Supplementary Figure S3) Switching to T2T from hg38 improved per-read sensitivity for host reads removal by 0.050-0.144 percentage point (all adjusted p-values < 0.001), and improved the overall performance, as measured by MCC, by 1.063-4.455 (all adjusted p-values < 0.001). If upgrading to T2T from YH, the increase in sensitivity were 0.3309-0.7238 percentage point (all adjusted p-values < 0.001), and the increase in MCC were 9.679-16.41 (all adjusted p-values < 0.001). (Table 1)

**Table 1.** Improvement of performance when switching to T2T.

|  | T2T compared to hg38 | T2T compared to YH |
| --- | --- | --- |

| Sequencer |  | Median of differences (pp.)<sup>*</sup> | Adjusted p-value | Median of differences (pp.) | Adjusted p-value |
| --- | --- | --- | --- | --- | --- |
| ILMN | Sensitivity | 0.053 | 1.15E-12 | 0.327 | 1.15E-12 |
|  | Specificity | -0.00628 | 1 | -0.00604 | 1 |
|  | MCC | 3.145 | 1.15E-12 | 14.838 | 1.15E-12 |
| MGI | Sensitivity | 0.144 | 7.82E-13 | 0.724 | 7.82E-13 |
|  | Specificity | 0.000514 | 1 | 0.000615 | 1 |
|  | MCC | 4.455 | 7.82E-13 | 16.413 | 7.82E-13 |
| ONT | Sensitivity | 0.050 | 1.50E-06 | 0.714 | 1.86E-09 |
|  | Specificity | -0.000682 | 1 | -0.0177 | 1 |
|  | MCC | 1.063 | 1.16E-04 | 9.679 | 1.64E-07 |

\* pp.: percentage point.

### Genomic location of reads exclusively mappable to T2T concentrated in newly added low-complexity regions

Reads that were exclusively mappable to the T2T genome, i.e., gold-standard human sequences that were unmappable to hg38 and YH (false negatives), were mostly located in the centromeric, acrocentric and telomeric regions using the T2T coordinates, most notably in the centromere of chromosome 1, 2, 3, 4, 5, 7, 9, 10, 16, and 20, and the short arms of chromosome 13, 14, 15, 21 and 22. This distribution is largely in agreement with the regions newly added to T2T.^9^ The other segments rich of T2T-exclusively mappable reads, e.g. those on chromosome 9, 16, and Y, may contain heterochromatic regions that are highly variable and largely genetically inactive, or sequences sharing similarity to those newly added to T2T (Figure 2A). We further characterized the false negative reads with hope to understand why they were not removed correctly. On one hand, true host reads that were unmappable to each genome showed a difference in their average guanine-cytosine (GC) content (Figure 2B). However, this difference was inconsistent across the sequencing platforms, and therefore was possibly the effect of sequencer bias, instead of reference genomes. On the other hand, >87.7% of the 75bp ILMN FN reads, >91.4% 50bp MGI and >26.2% of the long ONT FN reads were assigned by kraken2 as unclassified, cellular organisms and eukaryote (Figure 2C). This was largely due to low-complexity that results from the predominant (GGAAT)n repeats in human satellites^23^ (Figure 2D). The number of unclassified reads decreased for longer reads, as repetitive sequences were better mapped with increased read length. For short reads, T2T had much lower number of errors, possibly thanks to its revamping of the satellite regions.

**Figure 2.**
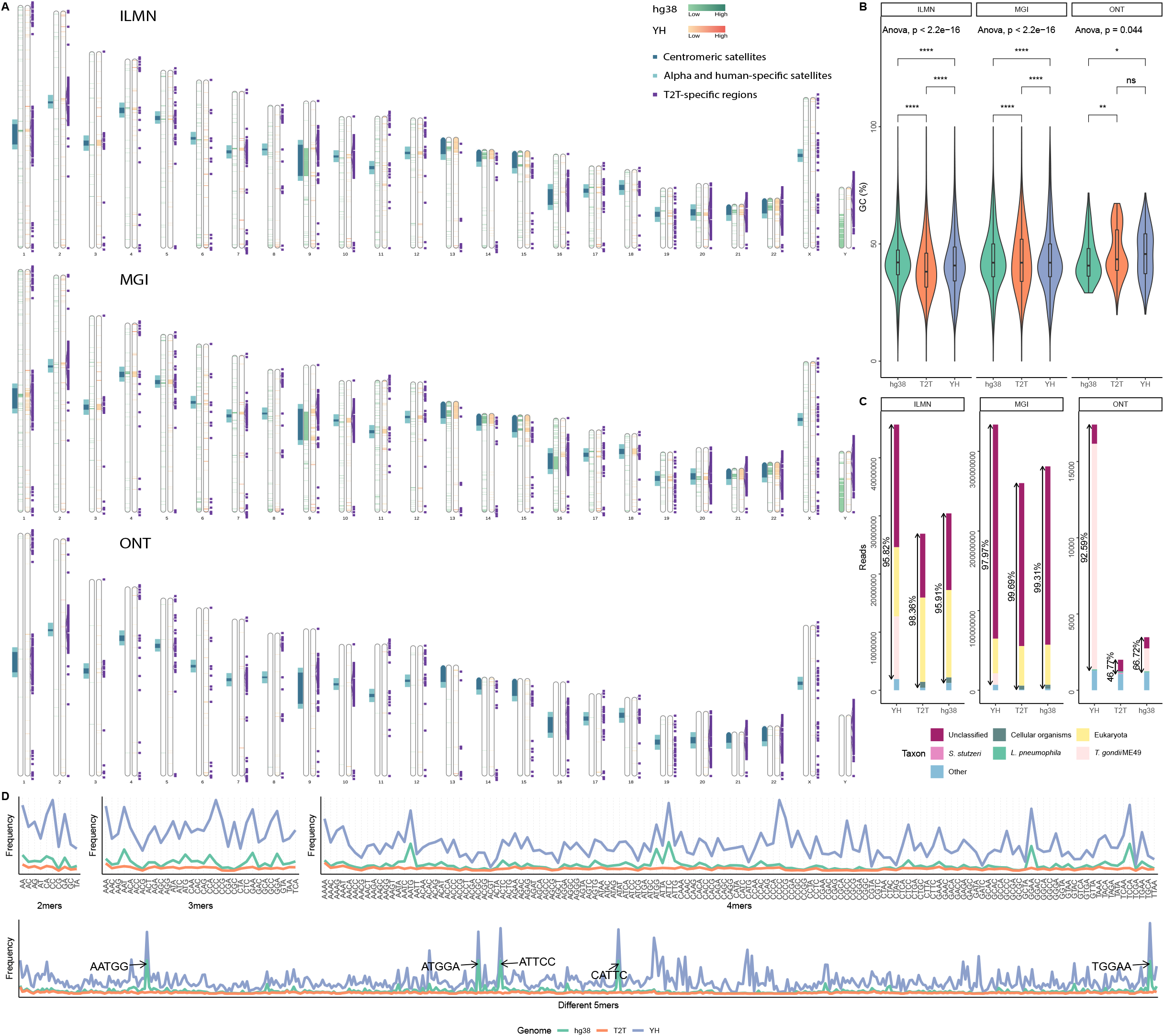
Characteristics of the uniquely mappable reads. **A** Ideogram of the T2T-CHM13 assembly, showing the genomic location of reads unmappable to the other reference genomes (for each chromosome, left column in green: hg38, right column in orange: YH); regions newly added therefore unique to the T2T assembly (purple); centromeric (deep blue) and alpha/human-specific (light blue) satellite arrays. **B** Distribution and comparison of GC content of reads unmappable to each reference genome. **C** Distribution of taxa falsely assigned to true host reads. Only the top 5 taxa are shown separately, the rest are tallied together. Percentages are the proportion made up by the top 5. **D** The k-mer spectrum analysis of false negative reads for each genome.

### T2T did not affect true positive calls but reduced false positive calls in microbe detection

By assigning microbial taxonomy to the non-host reads, including both true negative and false negative ones, we determined the impact of changes in host reads removal on the downstream microbe calling procedure. Noteworthily, true negatives in host reads removal constitute the pool for true positive calls in microbe detection, while false positive microbe calls arise from false positive host reads. We define the true positive microbe calls as reads whose gold standard label was non-host, and whose kraken2 classification was the organisms we spiked in the mock samples or culture-positive pathogenic organisms in the clinical samples. Across all sequencing platforms, three genomes showed very small differences (<0.21%) in true positive microbe calls (Figure 3A). Other commensal organisms were also detected in the clinical samples, for which we performed PCA analysis and found no difference between the results of all three genomes (Supplementary Figure S5). However, T2T greatly reduced false positive microbe calls in terms of both the number of false positive taxa and the false positive reads per taxon. In particular, we found that a large fraction of human reads was misclassified as *Toxoplasma gondii* ME49 (Figure 3B). This was likely due to the fact that the reference genome of *T. gondii* used in the kraken2 database (Accession number: GCA_000006565.2), which was sourced from RefSeq by default, contained contaminating human sequences, as reported previously^24^. Other important false positive taxa include *Staphylococcus aureus* and *Plasmodium vivax* (Figure 3B), both of which, like *T. gondii*, appeared in both short and long reads, suggesting contamination in the microbe’s reference genome. This was confirmed by genomic BLAST between the microbe and human reference genomes (Supplementary Table S4). Some high-ranking taxa, like Eimeriorina, Prioplasmida, and Streptomyces, only showed up for hg38 or YH in short reads, but were eliminated by T2T (Figure 3B). They were likely caused by imperfections in the previous human reference genomes, rather than microbe genomes. Indeed, in almost all cases, using T2T as reference genome greatly reduced false positives in microbe detection.

**Figure 3.**
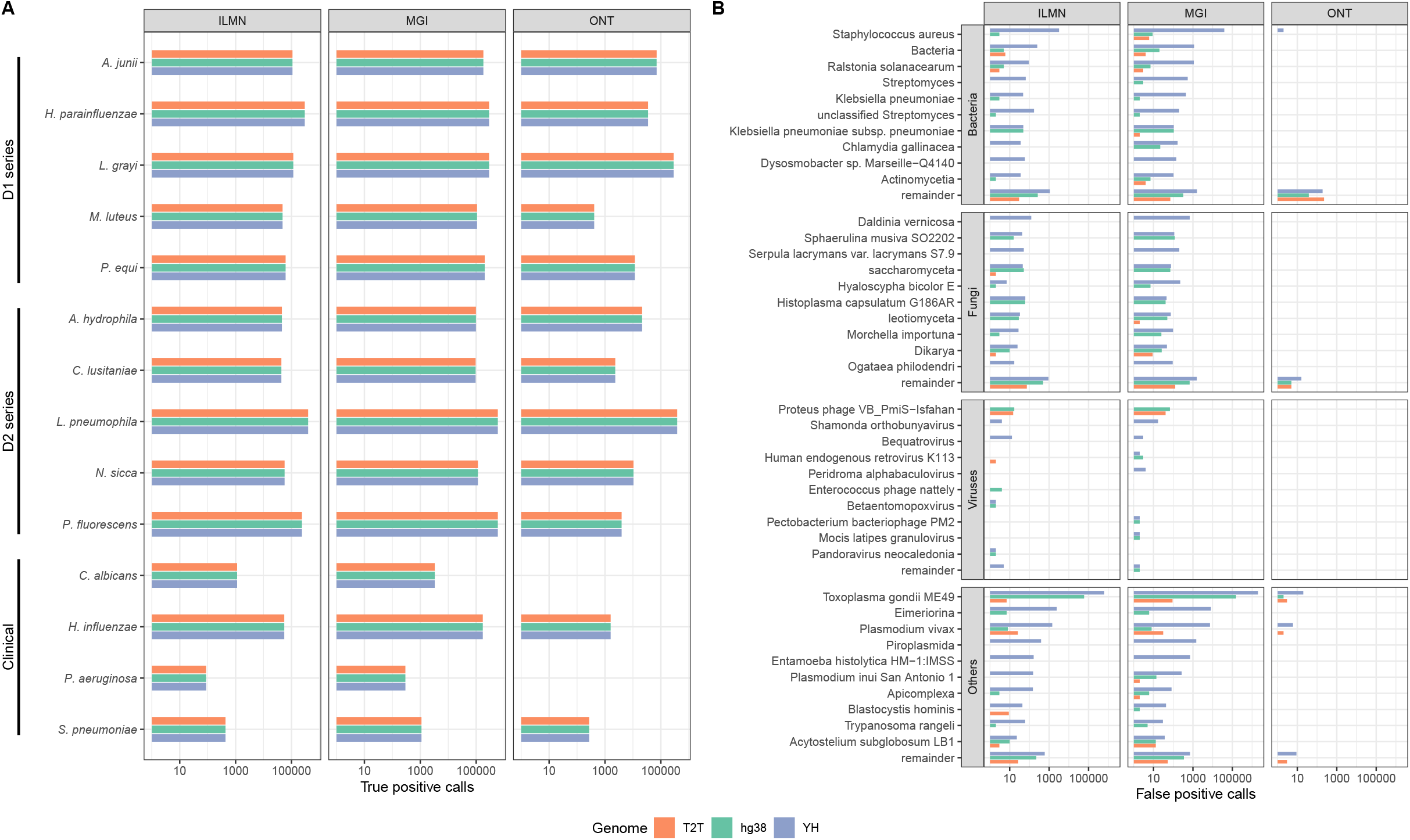
Impact of using T2T as reference genome on the results of microbe detection. **A** The true positive microbial sequence classification calls. A comparison between results of all the reference genomes is shown for both mock (D1 and D2 series) and clinical samples, across three sequencing platforms. **B** The false positive microbial sequence classification calls. The taxa reported by kraken2 are hereby categorized into bacteria, fungi, viruses and others, which mostly include amoebas and apicomplexans. Only the top 10 taxa are shown, all sequences classified as other taxa are tallied into the remainder group.

Interestingly, we also found a small number (<100 in total) of false positive reads for long reads that were classified to microbes we spiked in the mock samples. We suspected that they were unlikely to be truly of host origin, and the gold standard might have failed in these relatively rare cases. After examining the base-wise alignment of these reads (Supplementary Figure S6), we identified two possible causes: 1) ONT sequencer occasionally generated chimeric reads as the results of two DNA molecules passing the nanopore consecutively. These chimeric reads contained genuine human sequences and microbe sequences and thus were classified into both categories. 2) Sequencing errors could arise from pore destabilization and wobbling in nanopore sequencing.^25^ We detected this pattern in some of the ONT reads. It was likely this type of error that introduced segments of low-complexity sequences embedded in the genuine microbe reads and resulted in these reads being falsely mapped to the human genome. In this regard, these “false positive microbe reads” were actually true positives, and led to a slight overestimation of false positive rates. However, given the rarity of the errors, this would affect the conclusions to a minor extent at most.

## Discussion

With the arrival of the new T2T assembly, many areas of genomic studies have been revamped. Our results demonstrate that upgrading to the new reference genome improves the performance of host reads removal procedures commonly applied in metagenomic sequencing. We advocate the use of T2T in future studies and clinical applications where host reads removal is needed in metagenomic sequencing. For short-read sequencing, we suggest the use of bwa mem, while for long-read sequencing, we find no algorithm achieving consistently higher performance. It should be noted, however, that all software was tested with the default parameters. Different parameter settings will have an impact on the performance and may lead to different results. Fine-tuning is recommended, but it is beyond the scope of this study.

False negative and false positive calls in removing host reads have very different impact on downstream analysis. False negative means host reads getting passed the filter and entering the next stage in bioinformatics pipelines. This will increase the CPU-time needed for downstream analysis, and hence the overall turn-around time, which is critical in clinical applications. More importantly, it may result in false positive calls in microbe identification or other downstream analyses. In this regard, switching to the more sensitive T2T-based method will help to eliminate these artifacts. Given the inevitable imperfections in microbe reference databases due to its sheer scale, having T2T to maximize host reads removal will safeguard the microbe detection procedure against misassembled genomes in reference databases.

On the other hand, false positive in host reads removal leads to loss of microbe reads, and therefore decreasing the sensitivity of pathogen detection and hindering discovery of new pathogens. When preserving microbe reads are more of a concern, more specific approaches like STAT are recommended. Unfortunately, the current release of HRRT employs only a hg38-derived database and we invite the developers to consider upgrading the default reference genome to T2T. Moreover, we find that taxonomically misclassified reads are mainly the results of reference genome contaminations and sequencing abnormalities. This encourages future projects to put more efforts in removing contamination sequences in the reference databases, and to develop methods to reduce spurious reads.

In conclusion, using a diverse read set sequenced from real samples, we demonstrate that T2T brings improvement to correctly removing host reads in metagenomic experiments. The superiority of T2T over current reference genomes likely foretells its success to serve as the next prevailing reference genome in metagenomic sequencing, especially when host reads are overrepresenting.

## Materials and Methods

### Read set

In order to benchmark the performance of the bioinformatics methods in diverse biological settings, we sequenced both a set of mock community samples and a collection of clinical samples in this study. The mock community samples were prepared using human cell lines with gradient dilution of spiked-in microbes. Detailed setup of the samples was summarized in Supplementary Table S5. Clinical samples dataset (n=8) was desensitized, with all information that can identify the source of the samples removed. For mock samples, library construction and sequencing were conducted in parallel by 4 independent laboratories. The libraries were sequenced to 75bp single-end reads on Illumina NextSeq, and 50bp single-end reads on MGI MGISEQ-200 sequencer. In addition, library for long-read sequencing was constructed and sequenced on a Nanopore GridION platform. Following common practice, the raw sequencing data were preprocessed to remove adapter sequences, low-quality reads, short reads. The quality control was performed using fastp v0.23.2 for short reads, and Nanofilt v2.7.1 and NanoStat v1.5.0 for long reads. One ILMN library was excluded from the study due to insufficient sequencing yield.

### Host reads labelling

Reads that passed all quality controls were subjected to the host reads removal procedure. We applied a combination of reference genomes (T2T, hg38 and YH) and algorithms to the sequenced samples to give each read a label. For short-read sequencing, host reads removal were performed with two alignment-based algorithms: Bowtie2 v2.4.5 (in very fast and very sensitive mode) and bwa mem v0.7.17-r1188, along with two k-mer based algorithms: STAT (sra-human-scrubber or HRRT) v1.0.2021_05_05 and kraken2 v2.1.1. For long-read sequencing, minimap2 v2.17-r941 and winnowmap v2.03 were used as the alignment algorithms instead, with k-mer algorithms remaining the same. One exception was that host reads removal by STAT was only performed with hg38, because the HRRT distribution contained only a specially built database based on hg38, and we could not reproduce with the other two genomes. All software was run with default or preset (e.g. map-ont for minimap2) parameters. Each combination of the reference genomes and algorithms is hereafter referred to as a host-reads-removal “method”.

### Establishing the per-read gold standard

By each method, a read was given a label as whether it was of host origin (for kraken2, a TaxID assignment to any Chordata species was considered as a host label), but the true label of the read is unknown. Therefore, we compared all the labels given to each read to establish a *de facto* “ground truth”, or gold standard, by imposing the following criteria. A read was assigned a consensus label if concordant results were found for all methods. When the results were discordant, the reads were subjected to further examination. We used BLAST search results as the discriminating standard to resolve the discrepancy, because BLAST is an expensive yet very sensitive and more robust algorithm.^26^ Since the number of reads with discordant labels were too large to be all aligned by BLAST, we narrowed down the discrepancy by allowing a more tolerant standard for assigning consensus labels. If at least an alignment-based method and at least a k-mer-based method labelled a read as derived from host, we gave the host label to this read as the gold standard. The remaining reads with discordant labels were queried against the NCBI nr/nt database with BLAST (blast+ v2.12.0). If a hit to Chordata sequences were found with sufficiently high alignment quality (identity 90, coverage 90 for short reads and identity 70 for long reads), the read was considered truly of host origin, and otherwise non-host.

### Evaluating the performance

For each method, we compared the label it gives to each read against the gold standard, and classify the reads as true positive (TP), false positive (FP), true negative (TN), and false negative (FN), where positive is when a read is labeled human, and vice versa. We then calculated per-read sensitivity, specificity, and MCC for all methods, defined as:

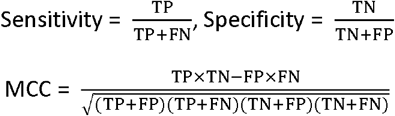

We included sensitivity and specificity as evaluation metrics because they are widely used and well-interpreted. Other commonly used metrics, e.g., in the CAMI benchmarking toolkit^27^, include the recall (or completeness) – precision (or purity) pair, and their harmonic mean, F1-score. These metrics involve only TP, but not TN in their calculation. Not including TN in assessing the performance of metagenomic profilers is justifiable by the fact that TN is hardly well-defined in this context. However, in samples with various host reads proportion, TP and TN are well-defined and can be overwhelmingly imbalanced. In high-human-read samples, where TP >> TN, recall (equivalent to sensitivity) and precision (equivalent to positive predictive value, or PPV) are inflated to 1 and lose discriminating power, and vice versa for specificity and negative predictive value (NPV). Therefore, we adopted MCC as an summary performance indicator, which has proven to be more balanced and reliable.^28,29^

When comparing the performance of the genomes, the results of the best performing software for each genome was used to represent the highest potential of a genome. When analyzing the unmappable reads, a read was considered falsely unmappable to a particular reference genome if the gold standard was positive yet none of the algorithms, when combined with that genome, gave a positive label. To further investigate the characteristics of these falsely unmappable reads, we examined the GC content, kmer frequency, false taxonomic classification and their true mapping location on the T2T genome, whenever applicable. The kmer frequency was calculated by jellyfish v2.3.0. We performed the PCA analysis with FactoMineR v2.5 and factoextra v1.0.7. We defined true positive microbe calls as those sequences whose gold-standard host label is negative and microbiological classification by kraken2 v2.1.1, with the default RefSeq database, is either spiked-in microbes in mock samples (Supplementary Table S5) or in accordance with the culture or FilmArray results in clinical samples (Supplementary Table S6). False negative microbe calls were defined as sequences with positive host labels in the gold standard but unremoved by a removal method and whose kraken classification does not belong to Chordata.

### Statistical analysis

To compare the sensitivity, specificity and MCC of all methods, we first tested the normality of the distribution of their difference using Shapiro-Wilk test and Q-Q plot (Supplementary Figure S7). As the assumption of normality was violated, we used non-parametric tests for comparison. Kruskal-Wallis test was used as the omnibus test, and Wilcoxon signed rank test (one-sided) with false discovery rate (FDR) adjustment was used as *post hoc* test for pairwise comparison. When comparing the performance of algorithms, the reference genome used was held invariable, to determine which algorithm served the best for a particular genome; when comparing the performance of the reference genomes, the best performing genome-algorithm combination was used to exclude the effect of the algorithms and compare the highest potential of a genome. Finally, to further investigate the unmappable host reads (FN reads) to each genome, we compared the GC contents of the unmappable reads by analysis of variance (ANOVA) and unpaired t tests (two-sided). All statistical analyses were conducted using R (v4.1.2). In all cases, a p-value <0.05 was considered statistically significant.

## Acknowledgement

The authors thank Genskey Co., Ltd. for providing computing resources to facilitate the benchmarking experiment of this study. The authors thank Lifeng Li and Donglai Liu for providing scientific insights and the sequences.

