## Supplementary Figure S1-S7 caption for "Telomere-to-Telomere Assembly Improves Host Reads Removal in Metagenomic High-Throughput Sequencing of Human Samples"

**Supplementary Figure S1. Sequencing statistics.** Count of raw reads, clean reads, mean length, q20, mean GC content, and duplication reads (Dup) for ILMN **A** and MGI **B**. Count of raw reads, clean reads, mean length, q10, and mean GC content for ONT **C**.

**Supplementary Figure S2. Workflow of collating host reads removal results of all methods.** Classification results from all combination of genomes and algorithms are collated together to establish the “gold standard”.

**Supplementary Figure S3. Comparison of sensitivity.** Sensitivity comparison across all reference genomes and relevant algorithms. Plots on the diagonal, first column and first row: distribution of per-sample sensitivity for all genomes, hg38 (green), T2T (orange), and YH (purple). Plots in the upper triangle: paired Wilcoxon (p-Wil) comparisons between each combination of reference genomes and algorithms. Numbers are median of differences between the pair, asterisks show the significance of adjusted p-values (no label: insignificant, *: <0.05, **: <0.01, ***: <0.001), and numbers in colors show the comparison for each genome. Plots in the lower triangle: scatterplot of sensitivity. Identity line (y=x) is shown in black, therefore points (each representing a sample) above it have higher sensitivity in y-axis than x-axis and vice versa. Abbreviations are vf: very fast, vs: very sensitive. Results of different sequencers were arranged in **I** for ILMN, **M** for MGI, and **O** for ONT.

**Supplementary Figure S4. Comparison of specificity.** Specificity comparison across all reference genomes and relevant algorithms. Plots on the diagonal, first column and first row: distribution of per-sample specificity for all genomes, hg38 (green), T2T (orange), and YH (purple). Plots in the upper triangle: paired Wilcoxon (p-Wil) comparisons between each combination of reference genomes and algorithms. Numbers are median of differences between the pair, asterisks show the significance of adjusted p-values (no label: insignificant, *: <0.05, **: <0.01, ***: <0.001), and numbers in colors show the comparison for each genome. Plots in the lower triangle: scatterplot of specificity. Identity line (y=x) is shown in black, therefore points (each representing a sample) above it have higher specificity in y-axis than x-axis and vice versa. Abbreviations are vf: very fast, vs: very sensitive. Results of different sequencers were arranged in **I** for ILMN, **M** for MGI, and **O** for ONT.

**Supplementary Figure S5. Principle Component Analysis of Commensal Organisms in the Clinical Samples.** The analysis of the composition of commensal organisms is based on classification by kraken2 after removing host reads with correspondent human reference genomes. Each point is marked by the combination of sample ID and the reference genome used. A good separation of samples is evident, but in general the results of all three genomes are identical and overlap with each other.

**Supplementary Figure S6. BLAST results of possibly chimeric reads and reads with sequencing errors.** Three examples are shown. **A** A chimeric read possibly raised from a human DNA molecule and a microbe molecule passing the nanopore in close succession. **B** A spurious microbe read with possibly a failure of the nanopore in middle of the sequencing process, resulting in a relatively low-complexity region that aligned to the human genome by chance. **C** A spurious microbe read with likely sequencing errors towards the end, possibly the results of failed nanopore or basecalling.

**Supplementary Figure S7. QQ plots of per-sample pairwise differences.** All numbers are compared to the value of T2T kraken2 as baseline. Deviation from the red line indicates violation of normality assumption of the distribution. **A-C** MCC for ILMN, MGI and ONT, respectively. **D-F** Sensitivity for ILMN, MGI and ONT, respectively. **G-I** Specificity for ILMN, MGI and ONT, respectively.
