## Supplementary figures and images for "Telomere-to-Telomere Assembly Improves Host Reads Removal in Metagenomic High-Throughput Sequencing of Human Samples"

### Supplementary Figure S1

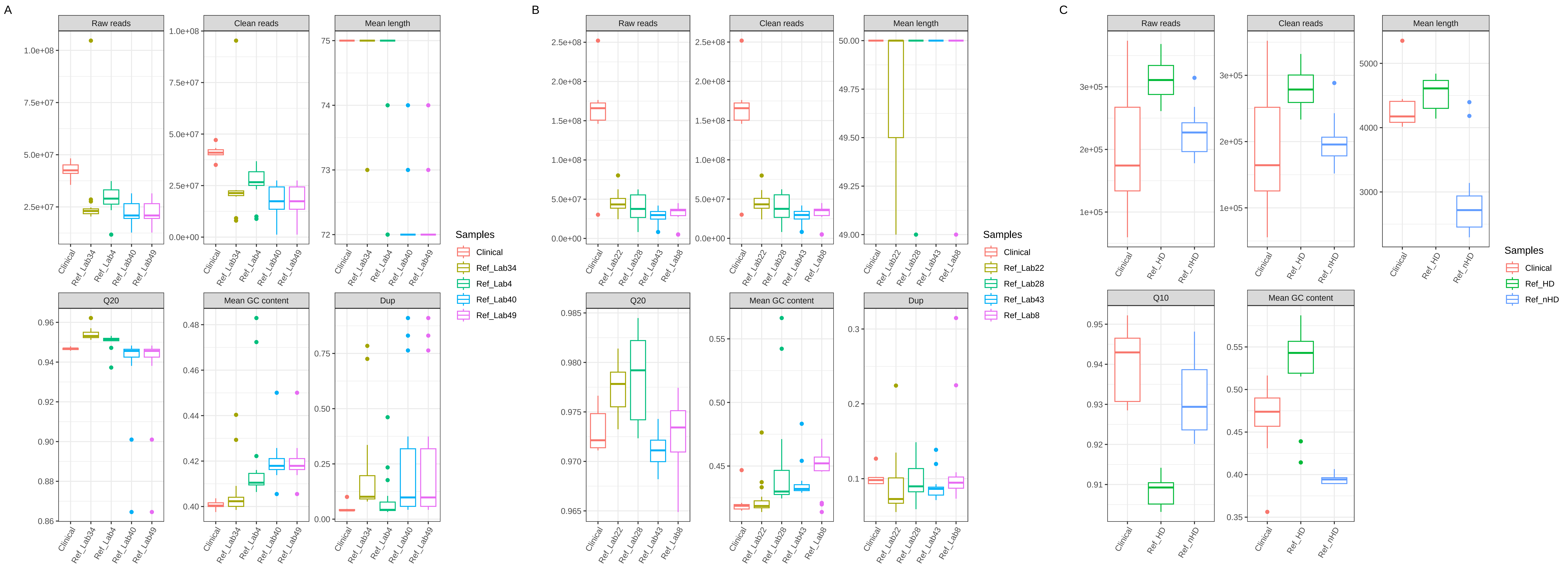

### Supplementary Figure S2

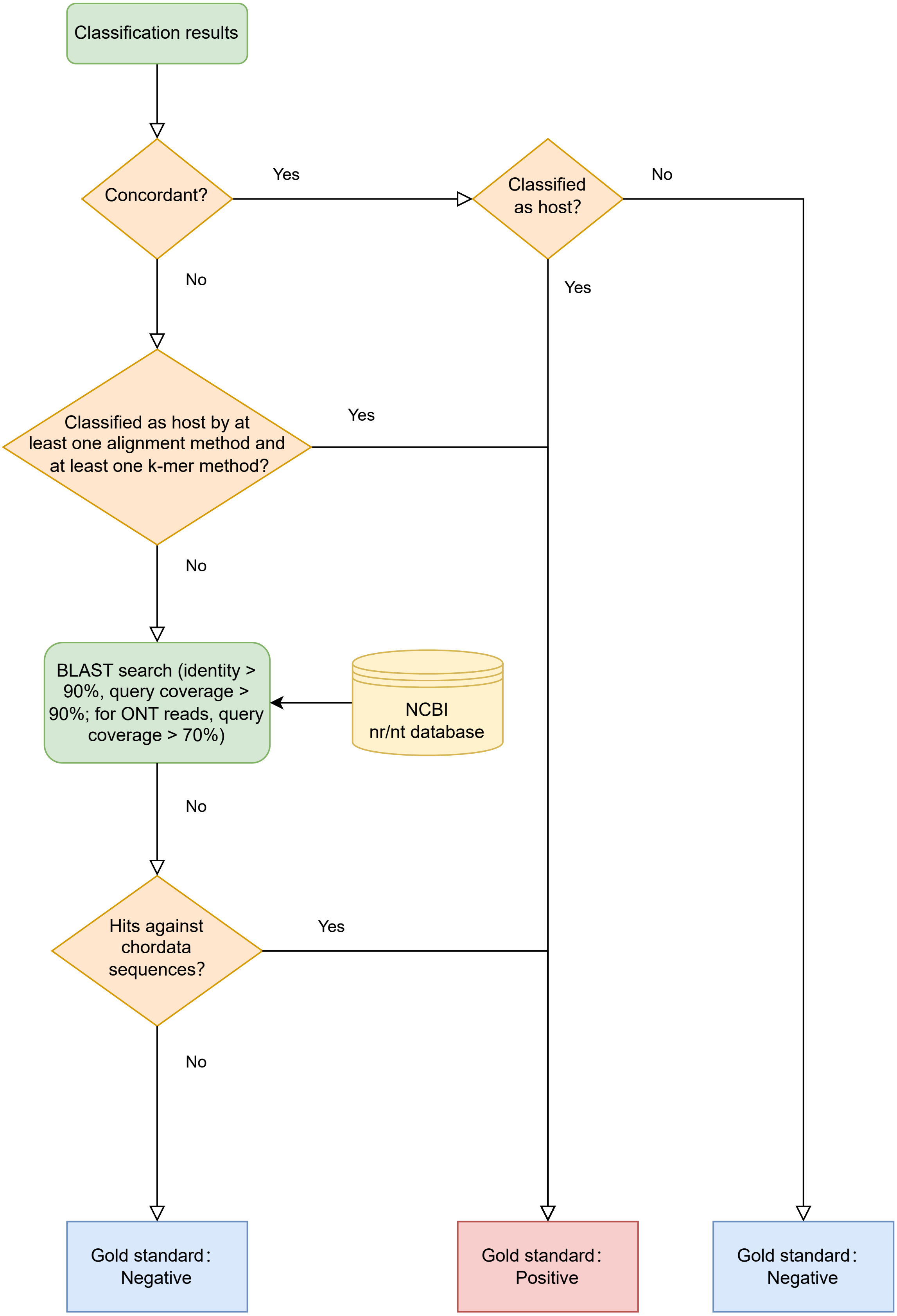

### Supplementary Figure S3

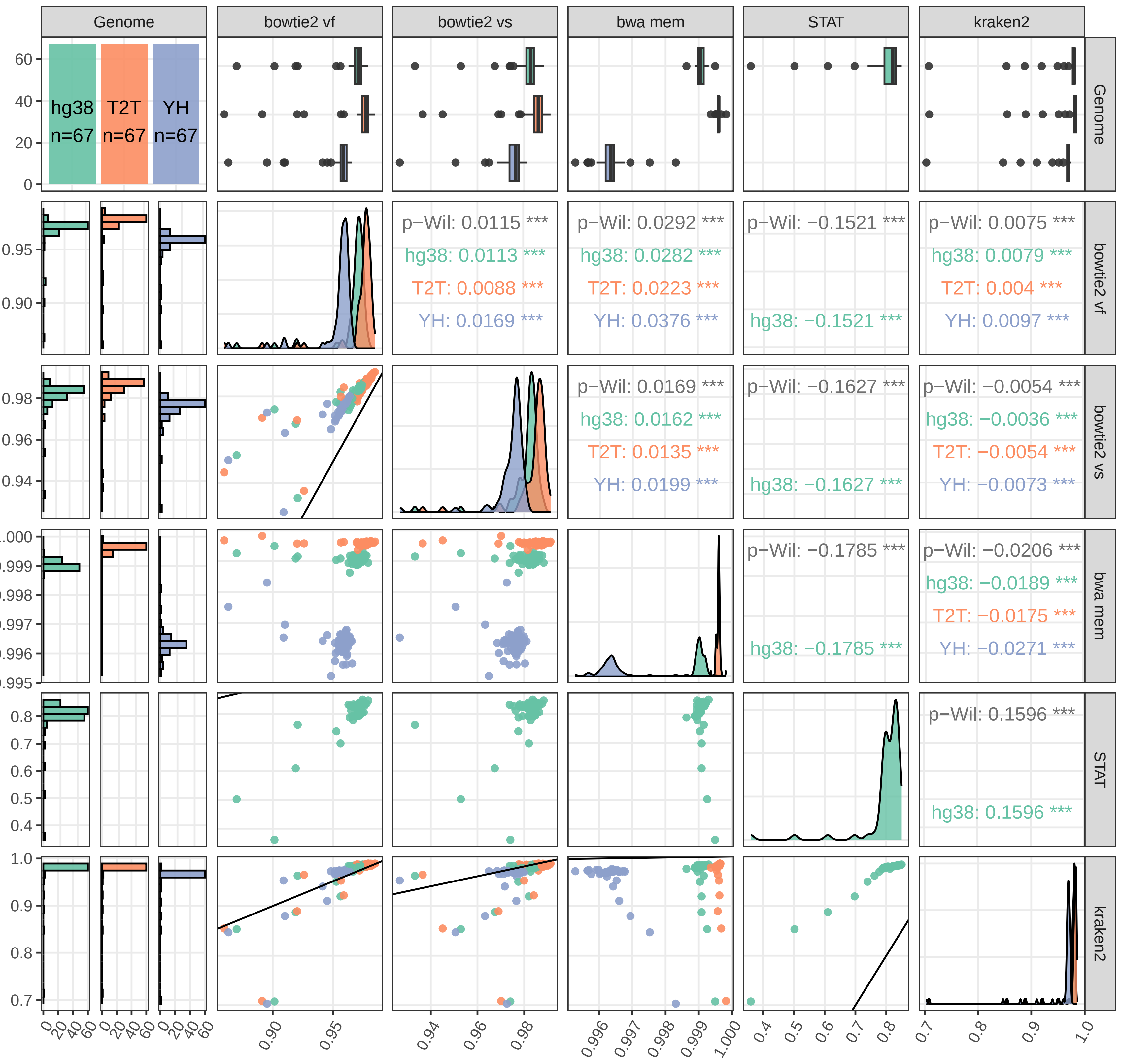

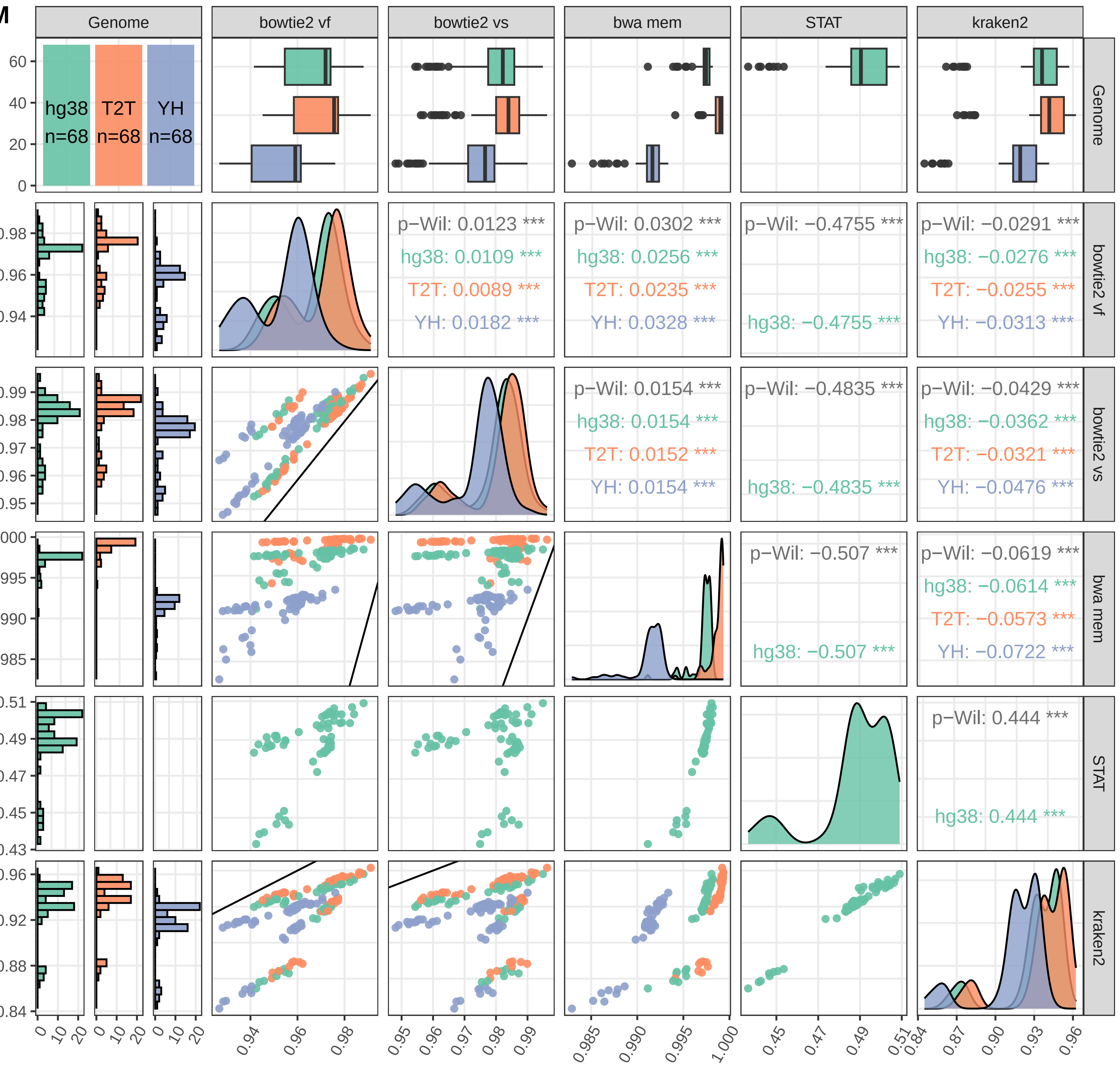

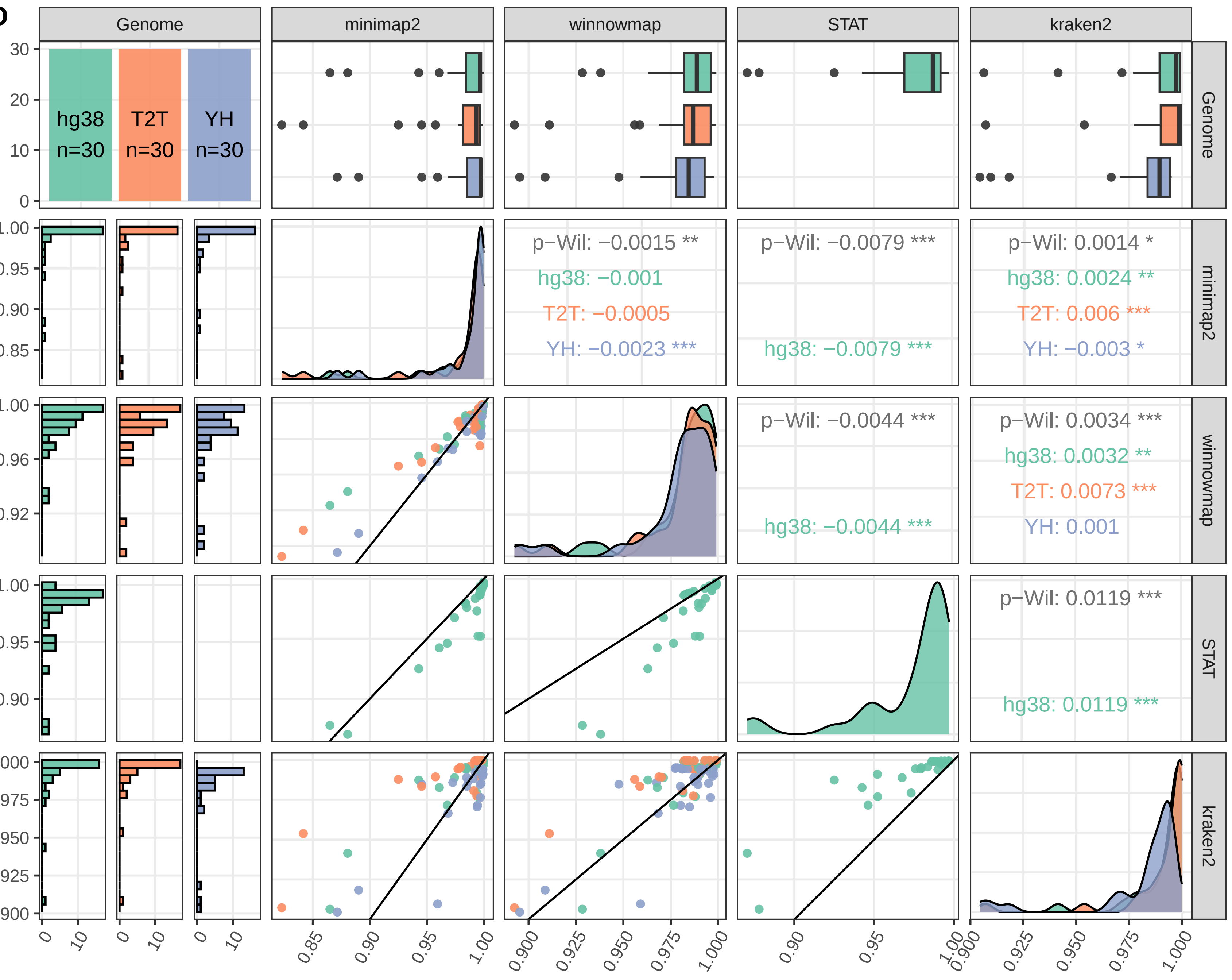

### Supplementary Figure S4

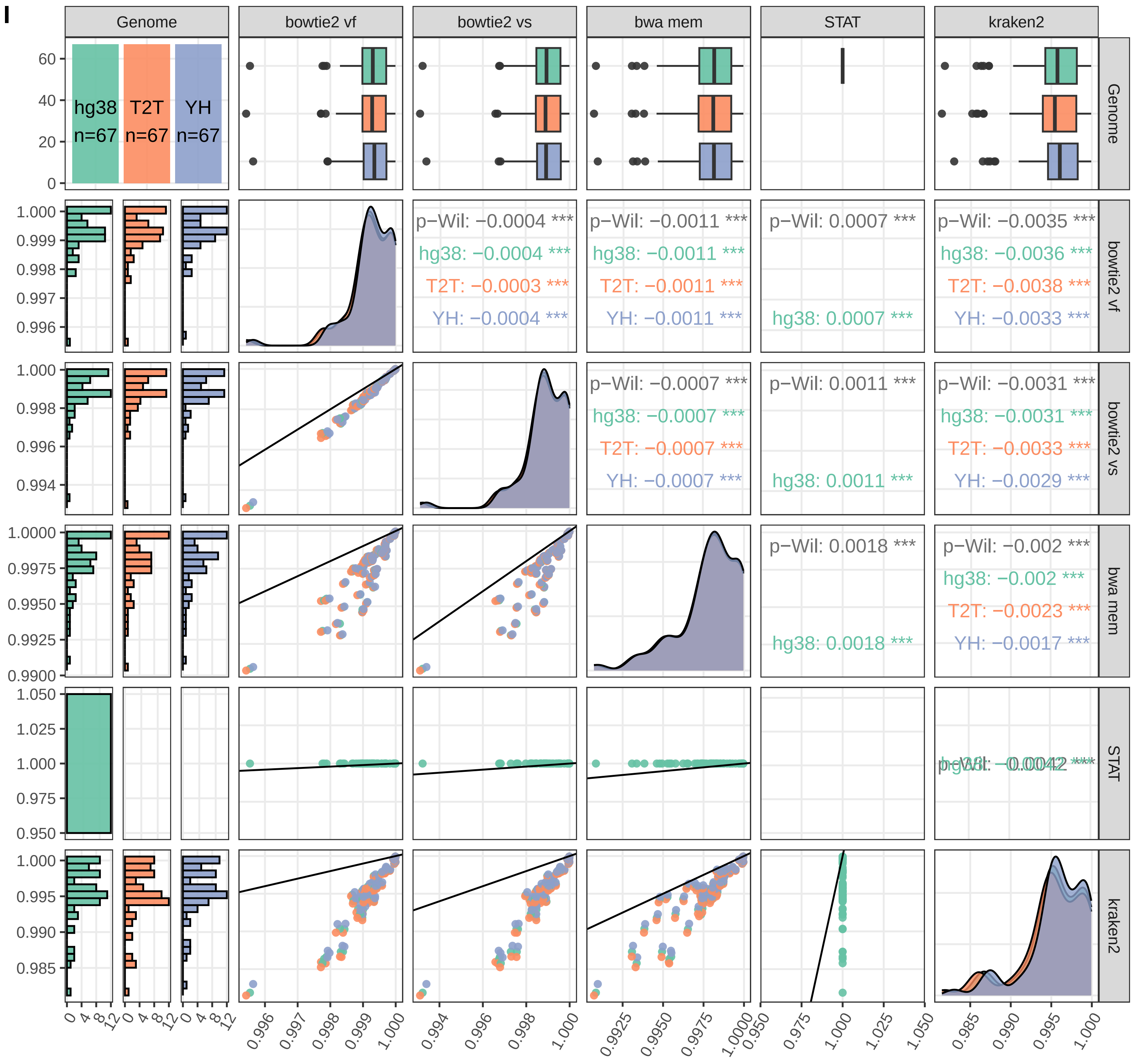

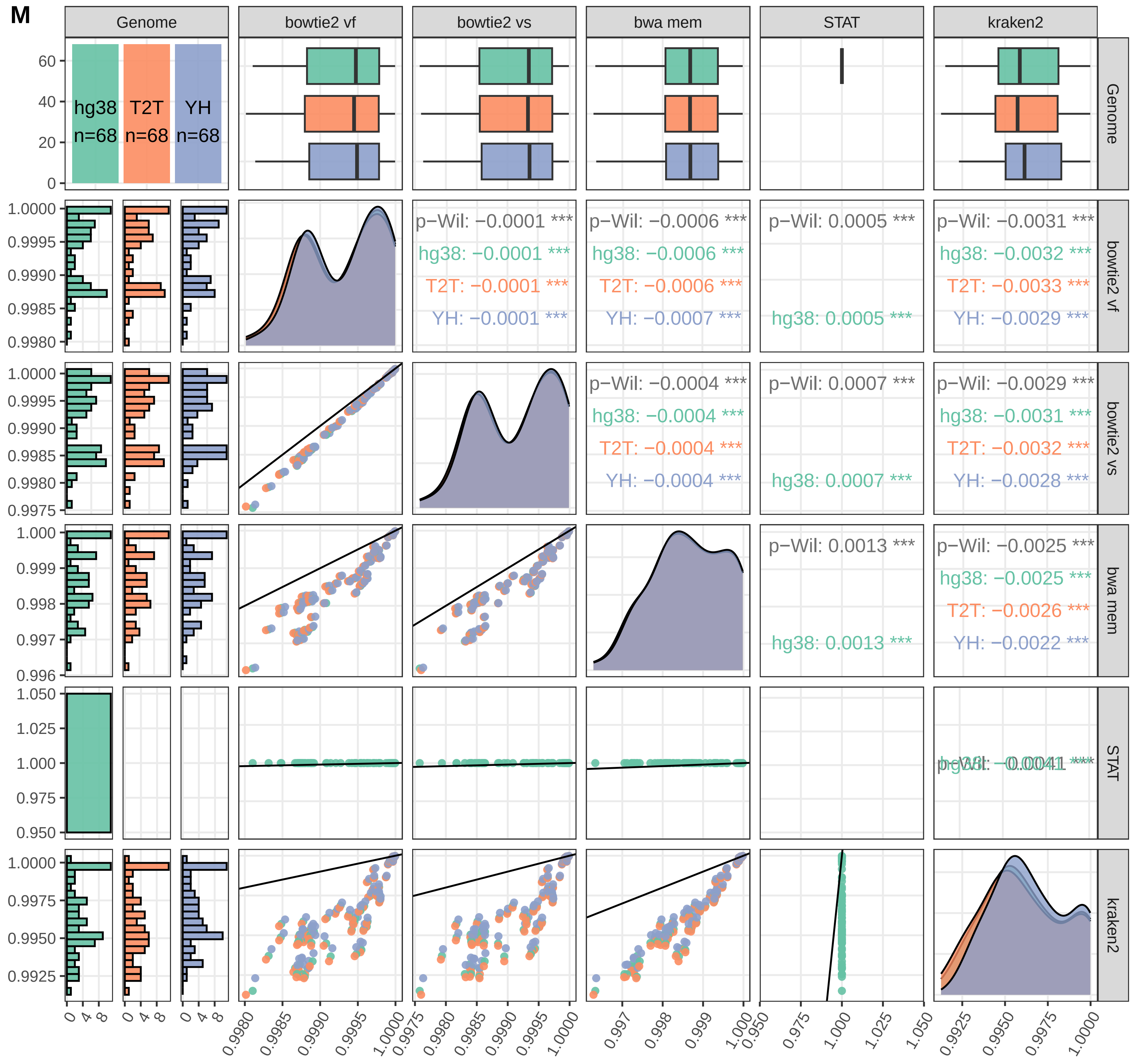

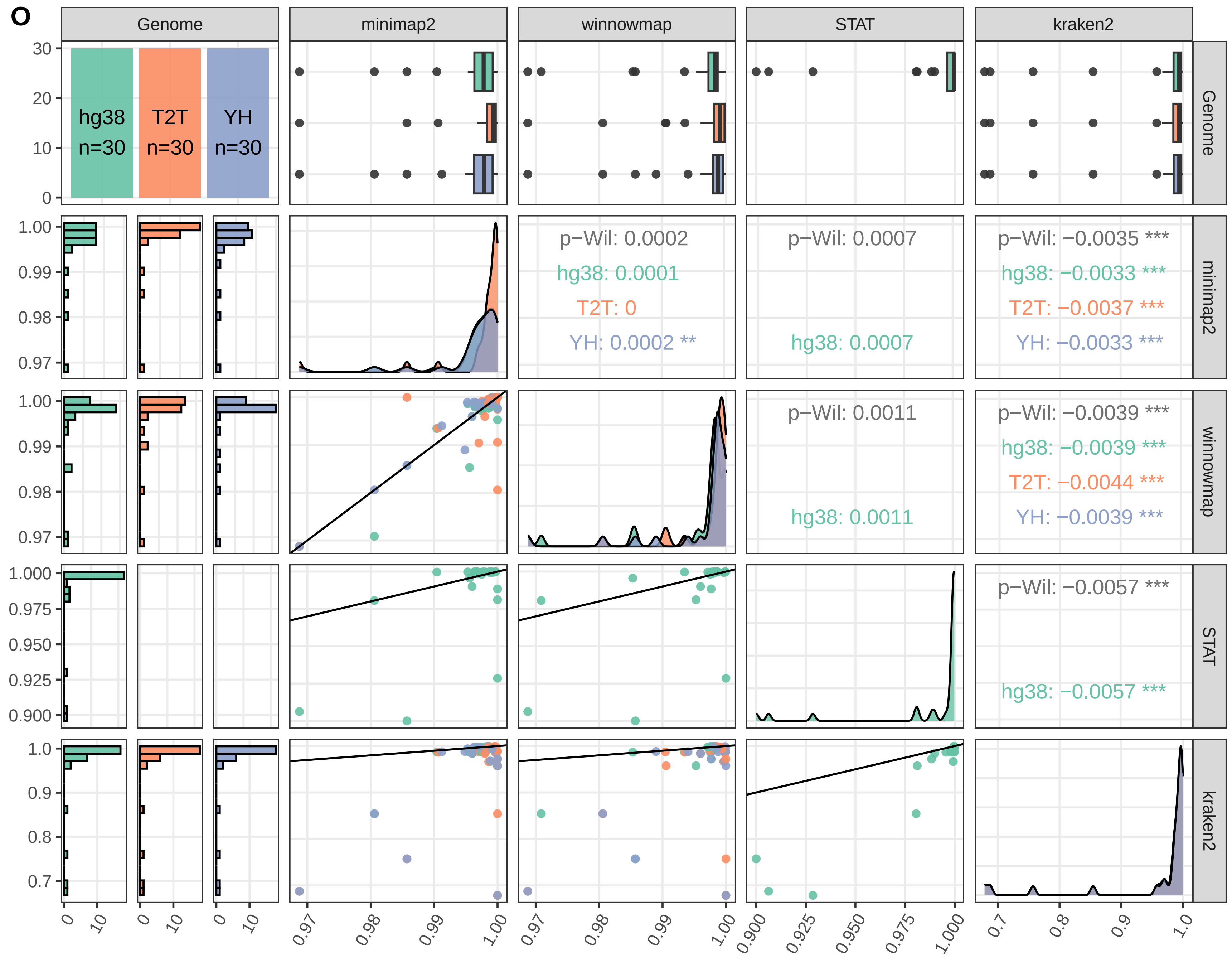

### Supplementary Figure S5

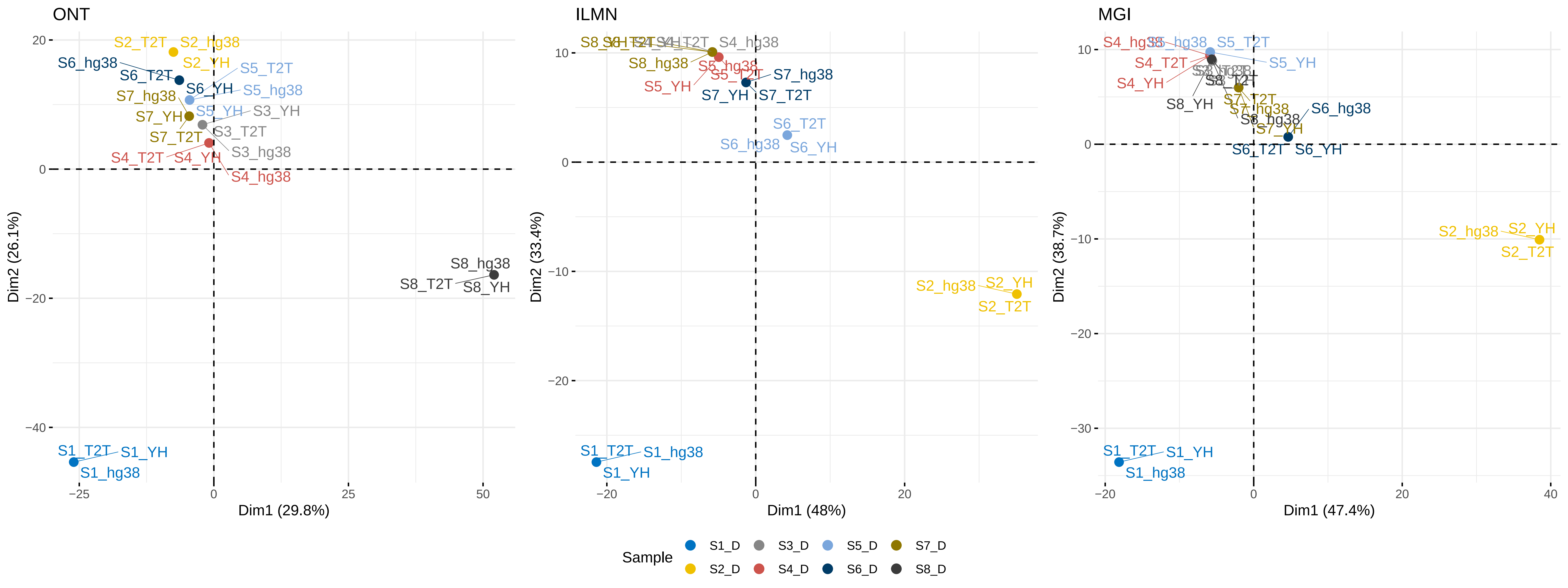

### Supplementary Figure S6

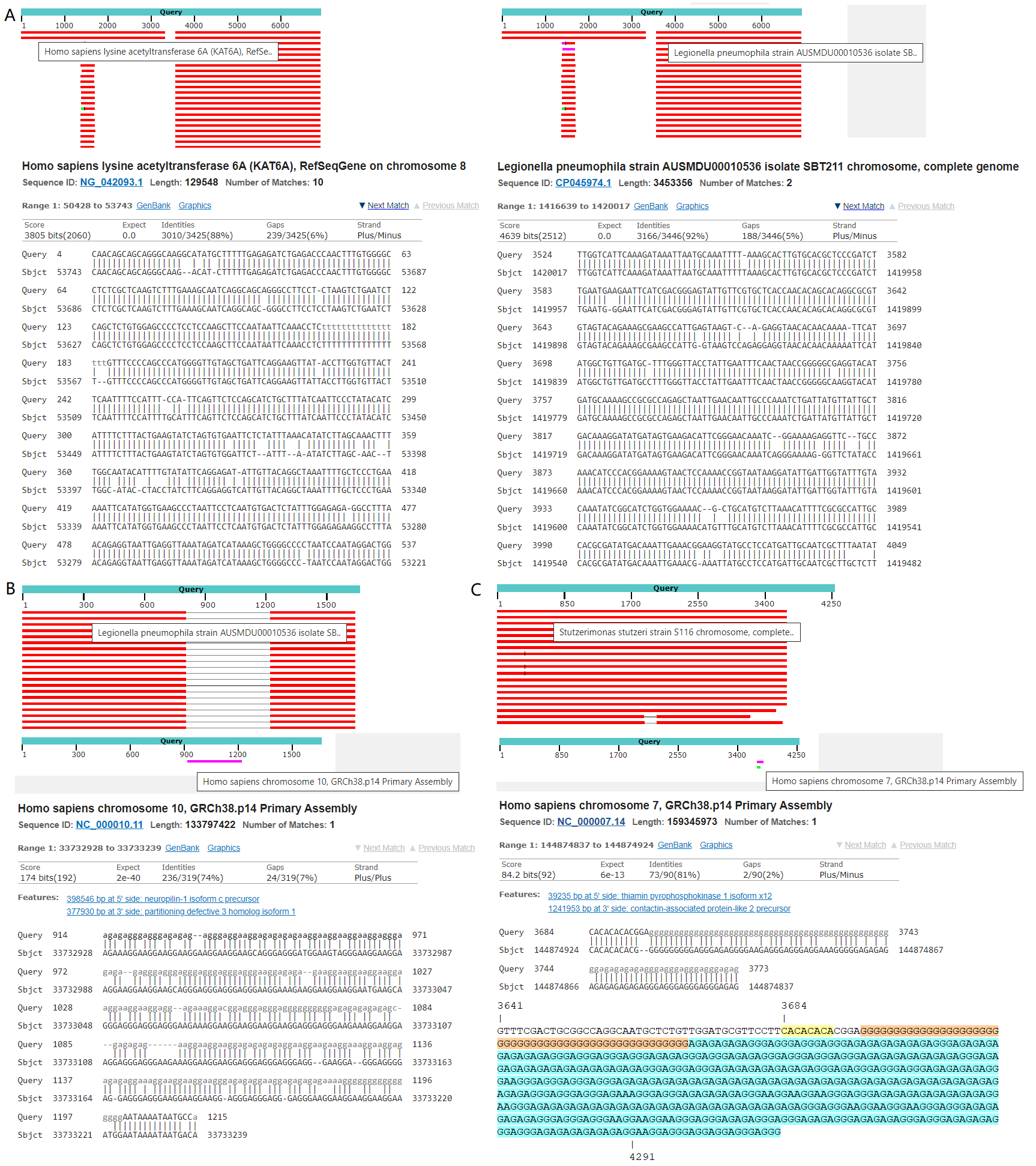

### Supplementary Figure S7

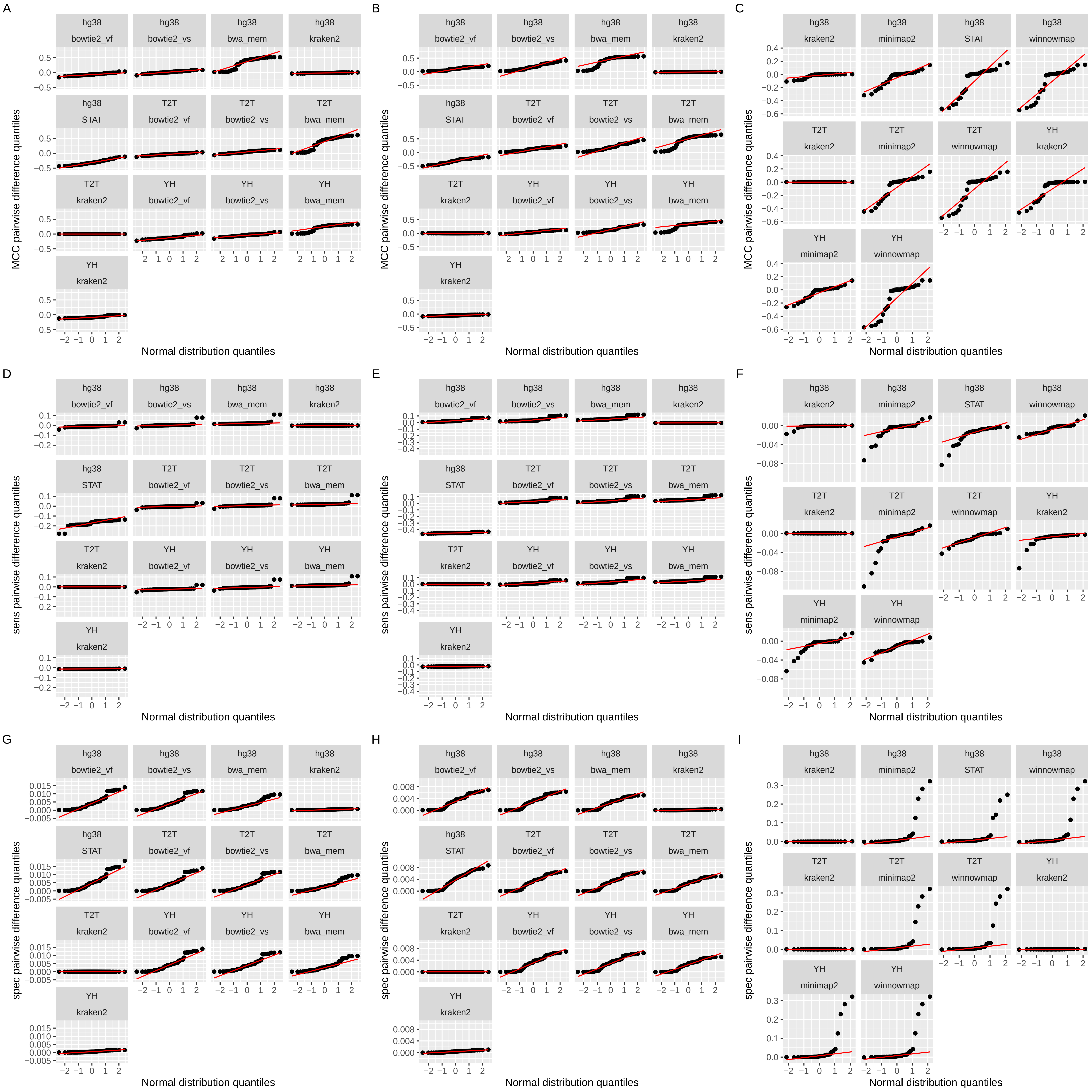
